# Acquired tick resistance in *Peromyscus leucopus* alters *Ixodes scapularis* infection

**DOI:** 10.1101/2025.04.22.650070

**Authors:** Elis A. Fisk, Cassie J. Leonard, Kristin L. Rosche, Elisabeth Ramirez-Zepp, Jeffrey R. Abbott, Jeb P. Owen, Dana K. Shaw

## Abstract

Ticks are obligate hematophagous parasites and pathogen vectors responsible for morbidity and mortality worldwide. *Ixodes scapularis* is a vector for at least seven pathogens relevant to human and animal health including the Lyme disease microbe, *Borrelia burgdorferi,* and the causative agent of anaplasmosis, *Anaplasma phagocytophilum*. Tick-host interactions are a driving influence on the maintenance of tick-borne pathogens in a population. Here, we report that repeated *I. scapularis* larval infestations on the wild host species *Peromyscus leucopus* leads to immune-mediated rejection of the tick, a phenomenon termed acquired tick resistance (ATR). We found that over 50% fewer larvae reached repletion and had decreased blood meal weights compared to larvae fed on naïve hosts. Additionally, mice exhibited increasingly severe inflammation at tick bite sites characterized by an influx of basophils, eosinophils, neutrophils, and T lymphocytes. Larvae fed on sensitized mice ingested higher quantities of host leukocytes when compared to ticks fed on naïve hosts, which rarely ingested nucleated cells. When challenged with *B. burgdorferi* or *A. phagocytophilum,* larvae fed on sensitized mice ingested more bacteria. Altogether, we demonstrate that reservoir host species develop ATR against larval *I. scapularis*, which reduces tick feeding success and affects pathogen ingestion by larvae. These results indicate that ATR could impact *Ixodes* population dynamics, prevalence of infected ticks, and pathogen circulation in the wild.

## INTRODUCTION

The incidence of tick-borne disease in the United States has been on the rise over the last decade, with 50,865 cases reported in 2019 increasing to 73,384 cases in 2022 (1). *Ixodes scapularis*, more commonly known as the deer tick, are capable of transmitting at least seven pathogens relevant to human health (2) including *Borrelia burgdorferi*, the causative agent of Lyme disease, and *Anaplasma phagocytophilum*, the causative agent of anaplasmosis. How the host and arthropod vector interact are a driving force influencing the maintenance of tick-borne pathogens in natural systems.

Some host species will develop protective immunity against ticks after repeated infestations, a phenomenon termed “acquired tick resistance” (ATR) (3). Tick salivary antigens elicit a T helper 2 (Th2) response (4), which stimulates IgE and/or IgG antibody production by B cells (5, 6). This IgE enters the bloodstream and arms the host’s circulating basophils and/or mast cells against tick antigens (6–10). Primed CD4+ memory T cells within the draining lymph nodes will then migrate to the skin. Upon subsequent tick bites, they secrete interleukin 3 (IL-3) which recruits basophils from the vasculature into the bite site (11). The recruited basophils along with resident mast cells will release histamine and other mediators which trigger local edema, itching, and epidermal hyperplasia (12–19). This type of immune response detrimentally impacts ticks by thwarting attachment success and reducing feeding weights, molting success, fecundity, and survival (3, 20–24).

Tick resistance has been well studied in laboratory animals, such as guinea pigs (*Cavia porcellus*), which develop ATR responses that reject over 80% of feeding ticks (20, 25, 26). For this reason, there is significant interest in developing translational strategies that will block tick feeding and pathogen transmission to humans and/or animals. In contrast, interactions with wild animal species are less well understood. The white-footed mouse, *Peromyscus leucopus*, is considered the reservoir host for *B. burgdorferi* and *A. phagocytophilum* and is often assumed to not mount ATR responses that rejects *Ixodes* ticks. However, some studies have reported varying degrees of resistance that appears to depend on life stage of the tick (27–30). While repeated nymphal infestations cause an increasing severe inflammatory response in *P. leucopus*, it does not hamper tick feeding (30). In contrast, repeated larval infestations result in lower feeding success, decreased weight, and reduced fecundity (27–29). Since nymph and adult life stages are primarily responsible for transmitting disease-causing microbes (31–33), these have been the main focus of ATR research (5, 34, 35). Larvae do not transmit *B. burgdorferi* and *A. phagocytophilum* (36, 37), but are the first life stage to become infected which makes them essential for pathogen maintenance in the population (31). The type of immunity elicited by *P. leucopus* to repeated larval infestation and whether this impacts pathogen acquisition is not known.

In this study, we demonstrate that white footed mice mount ATR responses against *I. scapularis* larvae, which impacts tick feeding and pathogen ingestion. We found that previously infested hosts exhibited severe inflammation at sites of larval attachment predominated by basophils, eosinophils, and neutrophils with epidermal ulceration and hyperplasia. In contrast, tick-naïve mice exhibited only mild to moderate inflammation predominated by macrophages, eosinophils, and neutrophils. The severe inflammation observed in sensitized mice correlated with a significant reduction in larval feeding success, fewer larvae reaching repletion, and decreased blood meal volumes. Additionally, the quantity of leukocytes ingested by feeding larvae increased proportionately with the number of prior infestations experienced by the host. We found that larvae ingested a greater number of *B. burgdorferi* from sensitized male mice, whereas *A. phagocytophilum* ingestion was enhanced with sensitized female mice. Altogether, we demonstrate that wild host species develop ATR against larval *I. scapularis,* which hampers tick feeding and alters pathogen acquisition.

## RESULTS

### Larvae feed less successfully on tick-sensitized P. leucopus

Both larval and nymph life stages are important for the cycle and maintenance of pathogens in a wild population. However, little is known about wild hosts responses to larval infestation. We therefore sought to quantify how previous tick exposure in *P. leucopus* influenced larval feeding success. *P. leucopus* mice were infested with 100 *I. scapularis* larvae one to four times (Fig 1A) and ticks were allowed to feed to repletion over 7 days. For mice infested more than once, a two-week waiting period was observed between infestations to allow an adaptive immune response to develop. After each infestation, the proportion of larvae feeding to repletion and the replete larval weight were quantified. During primary infestations, male mice supported an average of 19.1% (17.2%-21.0%) of larvae, whereas sensitized mice only supported 5.8% (4.7%-6.8%), representing a 70% reduction in feeding success (Fig 1B). Similar decreases in feeding success were observed in larvae fed on female mice, with up to 58% fewer larvae reaching repletion on sensitized mice (descriptive statistics summarized in Tables 1-2). We also noted that larvae fed more successfully on naïve males when compared to naïve females. However, following at least one tick sensitization, both sexes supported similar numbers of larvae.

**Figure 1.**
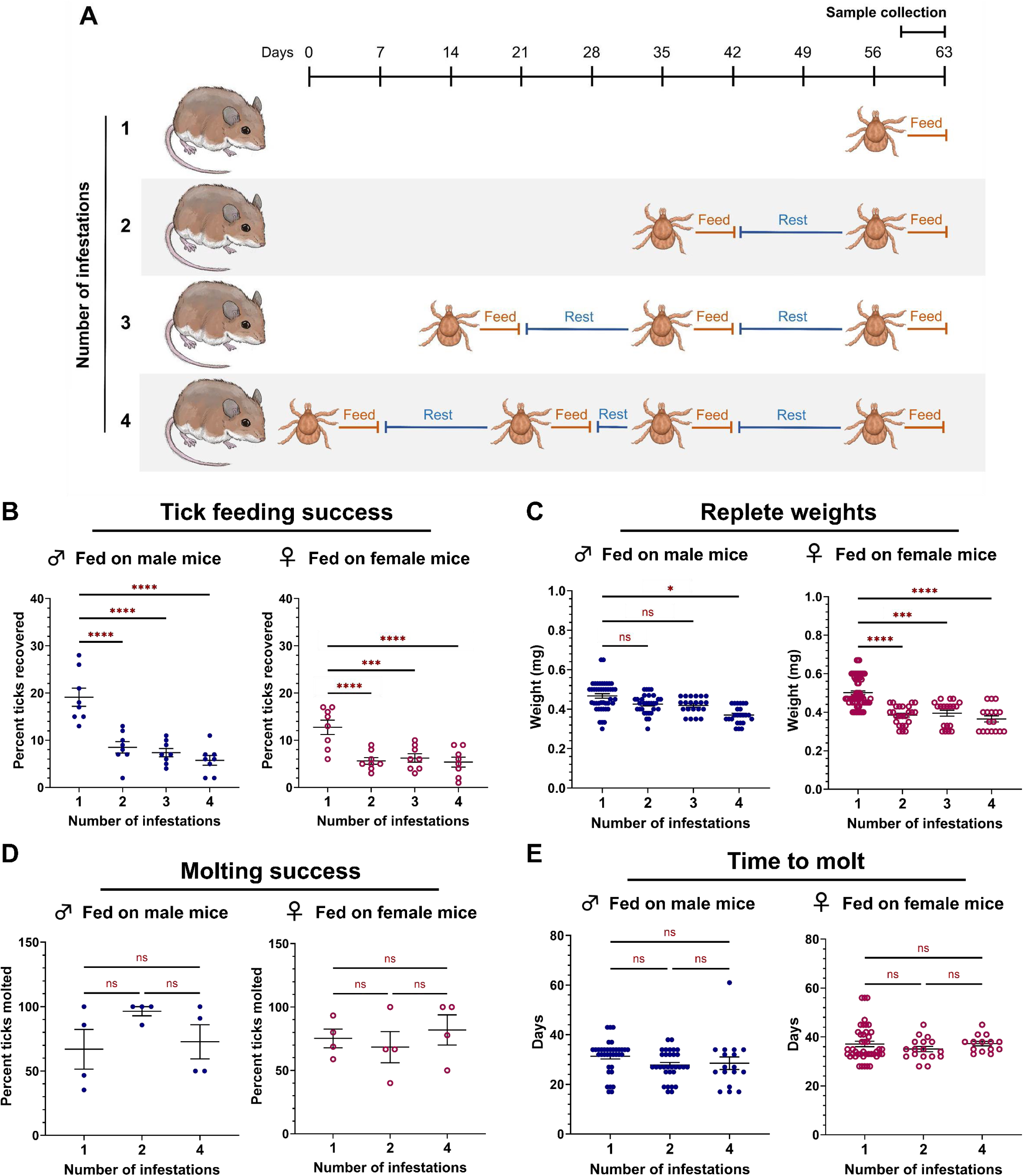
Reduced *I. scapularis* larvae feeding success on previously sensitized *P*. *leucopus*. (A) Schematic depicting the infestation schedule. Each larva is representative of one infestation event in which 100 larvae were manually placed on an anesthetized mouse. Larvae were allowed 7 days to feed to repletion. Animals used for histopathology and leukocyte analysis followed a similar infestation schedule with the final sample collection occurring 72 hours post-attachment. (B) Percent feeding success, (C) repletion weights, (D) molting success, and (E) time to molt were quantified for all conditions. Each data point represents either the proportion of larvae feeding to repletion on a single animal (B,D) or a single replete larva (C,E). A negative binomial generalized linear mixed effect model was used for statistical analysis of larval feeding success and time taken to molt. Larval replete weight was analyzed using a linear mixed effects regression model. Molting success was analyzed using a generalized linear mixed-effects model. **p < 0.01; ***p < 0.001, ****p<0.0001. ns = not significant.

To quantify blood meal volume, replete larvae were weighed. Larvae fed on naïve female mice had significantly larger blood meal volumes (0.502 mg ± 0.009) compared to those fed on sensitized mice (Fig 1C). Larvae fed on mice during the fourth infestation ingested 0.365 mg (± 0.016), representing a 27% reduction in blood meal volume. For larvae fed on male mice, an 18% drop in blood meal volume was observed between the first infestation (0.456 mg ± 0.010) and fourth infestation (0.376 ± 0.009). Although ticks fed on naïve mice had larger bloodmeals, there were no differences in the proportion of larvae molting successfully (Fig 1D) or the time taken to molt (Fig 1E). Bloodmeal volumes and molting success were also comparable between ticks fed on either male or female mice. Taken together, these data demonstrate that *I. scapularis* larvae feed less successfully on sensitized *P. leucopus* and acquire less blood, which is consistent with previous reports (27–29).

### Tick sensitization is associated with severe inflammation at the larval attachment site

We next histologically characterized the host’s skin response to larval attachment. *P. leucopus* mice were infested with larvae one to four times (Fig 1A). During the final round of infestations, mice were euthanized at 72 hours and punch biopsies were taken at larval attachment sites. When compared to naïve *P. leucopus* (Fig 2A-B), mice infested once showed a mild to moderate focal inflammatory response at the bite site (Fig 2C-D). Although the epidermis adjacent to larval attachment sites was mildly hyperplastic (25.2 µm ± 2.0 for males, 36.8 ± 6.5 for females), epidermal thickness did not differ significantly from uninfested mice (12.4 µm ± 1.2 for males, 17.2 ± 0.7 for females). In contrast, mice infested four times showed widespread and severe inflammation at the bite site (Fig 2E-F). The epidermis abutting larval mouthparts was eroded or ulcerated with serocellular crusting and the intact epidermis at the bite site exhibited significant hyperplasia (57.7 µm ± 15.7 for males, 88.0 ± 15.2 for females) compared to uninfested skin (Fig 2E-F; Supplementary Fig 1; Tables 1-3). These findings demonstrate that the inflammatory response to larval attachment is more severe in *P. leucopus* that were previously sensitized.

**Figure 2.**
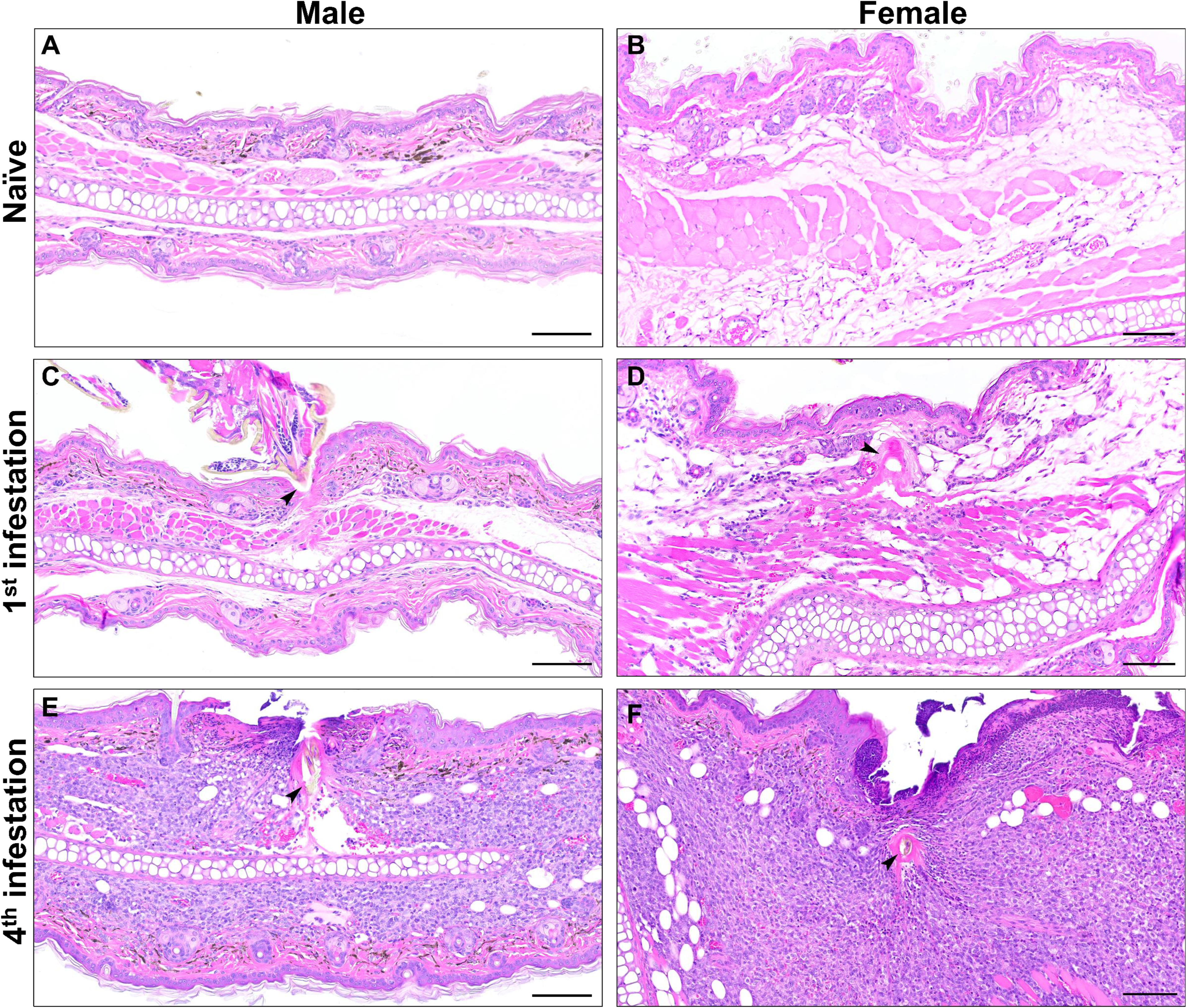
Increasing inflammation at larval attachment sites with previous tick exposure. Histological examination of male and female mouse biopsies from (A-B) naïve pinna, and (C-F) larval attachment sites from (C-D) a primary infestation and (E-F) a quaternary infestation. (C-F) Lymphocyte, macrophage, and eosinophil infiltrate in the dermis and centered around the hypostome and cement (arrow). (E-F) Ulcerated epidermis adjacent to the embedded hypostome (arrow) with serocellular crusting. H&E stain. Bar = 100 μm.

### Larval bite sites reveal an influx of inflammatory cells with repeated exposure

We next sought to characterize the immune cell population mediating the inflammatory response at larval bite sites. Skin samples were taken from mice infested either one or four times for immunohistochemistry, differential staining, and quantitative reverse transcriptase PCR (qRT-PCR). Mast cells and eosinophils were visualized with toluidine blue and luna stains, respectively. Neutrophils (MPO), basophils (MCPT8), T lymphocytes (CD3), and macrophages (IBA1) were visualized by immunohistochemistry with antibodies against specific cell type markers. We found that mice infested once showed a small number of eosinophils (Fig 3D), neutrophils (Fig 3G), and macrophages (Fig. 4G) around the larval bite site. In contrast, mice infested four times showed a robust inflammatory infiltrate composed of eosinophils (Fig 3E), neutrophils (Fig 3H), basophils (Fig 4B), and a moderate number of T lymphocytes (Fig 4E) and macrophages (Fig 4H). Sensitized mice showed neutrophils and basophils clustered around and extending into the cement cone of larval ticks (Fig 3H, Fig 4B).

**Figure 3.**
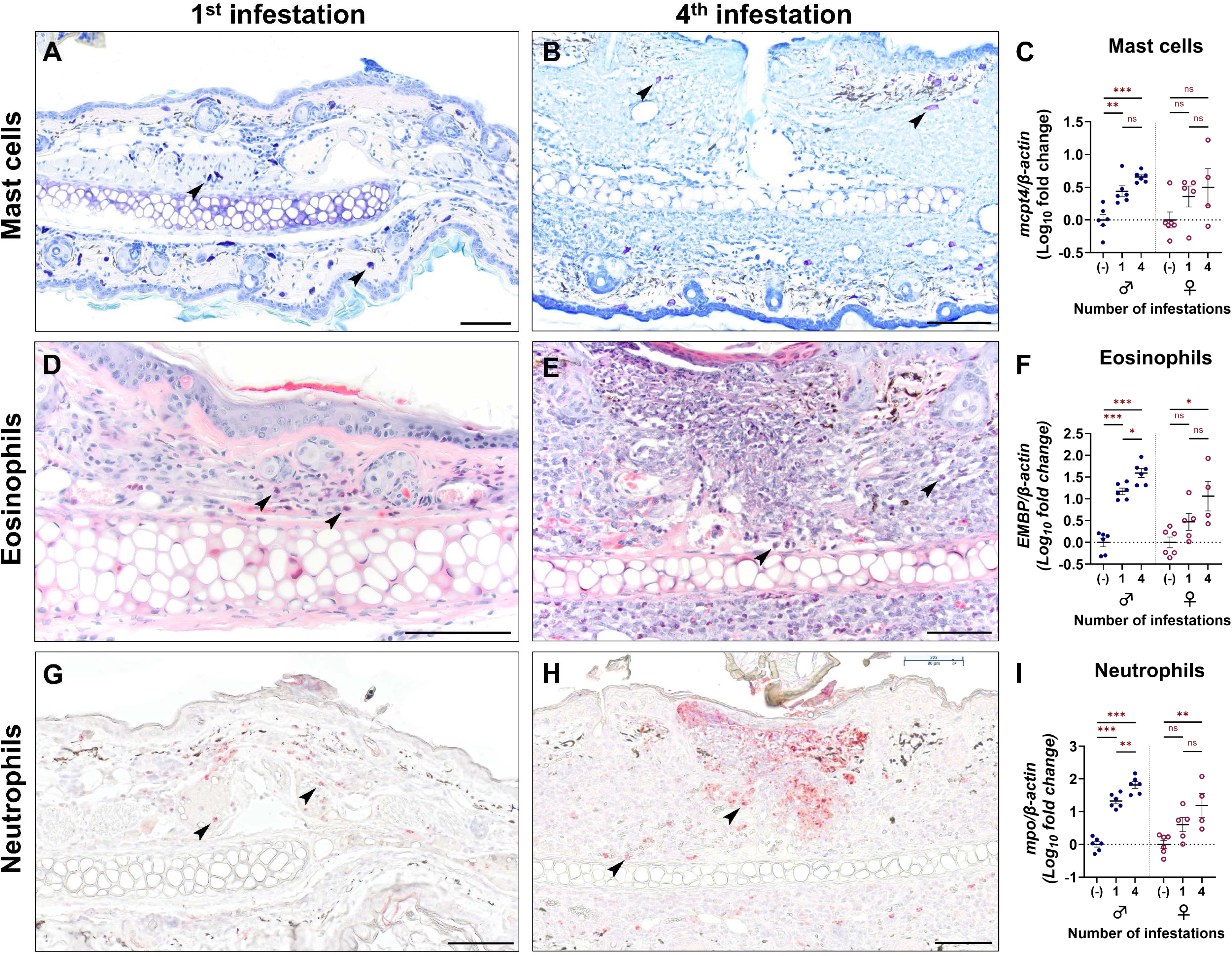
Eosinophils and neutrophils infiltrate larval bite sites with serial infestation. Leukocyte characterization at larval attachment sites using (A-B, D-E) special stains, (G-H) immunohistochemistry in primary and quaternary infestations and (C, F, I) gene expression in pinnal biopsies from naïve mice or at larval attachment sites from male and female mice. (A-B) Mast cells are visualized with toluidine blue stain (arrows). (C) qRT-PCR quantification of mast cell-specific transcript mast cell protease 4 (*mcpt4*). (D-E) Eosinophils are visualized by Luna stain (arrows). (F) qRT-PCR quantification of eosinophil-specific eosinophil major basic protein (*EMPB*). (G-H) Neutrophils are visualized by immunohistochemistry against Myeloperoxidase (arrows). (I) Expression of neutrophil-specific myeloperoxidase (*mpo*). All qRT-PCR data points represent individual bite sites or biopsy samples collected from the pinnae of uninfested mice. Leukocyte marker quantification was analyzed using a linear regression model. Bars = 100 μm. *p < 0.05; **p < 0.01; ***p < 0.001. ns = not significant.

**Figure 4.**
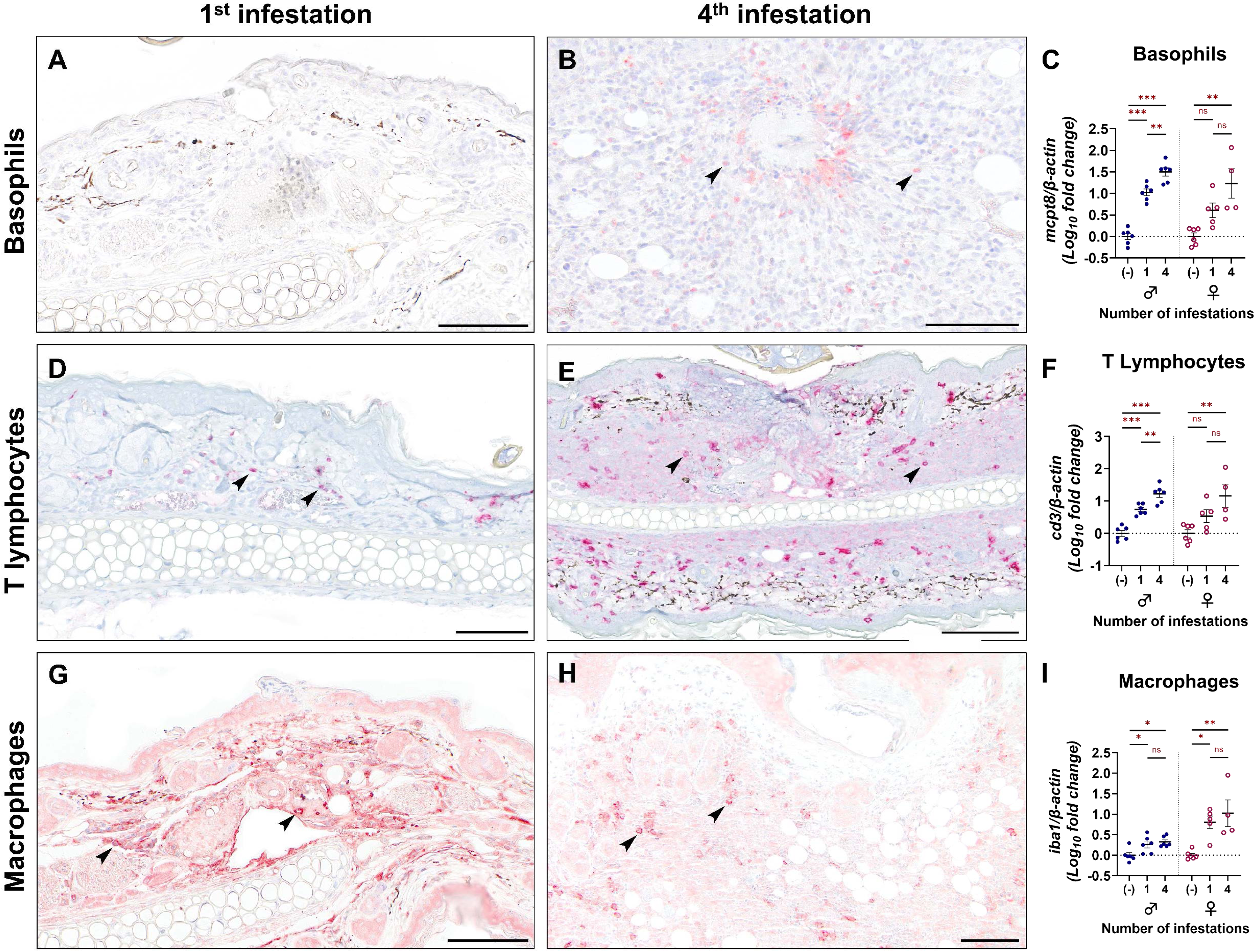
Serial infestation with larval *I. scapularis* elicits basophil and T lymphocyte infiltrate. Leukocyte infiltrate at larval attachment sites using immunohistochemistry in primary and quaternary infestations (A-B, D-E, G-H) and gene expression (C, F, I) in pinnal biopsies from naïve mice or at larval attachment sites from male and female mice. (A-B) Basophils were visualized by immunohistochemistry against Mcpt8 (mast cell protease 8) (arrows). (C) qRT-PCR quantification of basophil-specific transcript *mcpt8*. (D-E) T lymphocytes were visualized by immunohistochemistry against CD3 (cluster of differentiation 3) (arrows). (F) qRT-PCR quantification of T lymphocyte-specific transcript *cd3*. (G-H) Macrophages were visualized by immunohistochemistry against ionized calcium-binding adaptor molecule 1 (IBA1) (arrows). (F) qRT-PCR quantification of macrophage-specific transcript *iba1*. All qRT-PCR data points represent individual bite sites or biopsy samples collected from the pinnae of uninfested mice. Leukocyte marker quantification was analyzed using a linear regression model. Bars = 100 μm. *p < 0.05; **p < 0.01; ***p < 0.001. ns = not significant.

Inflammatory cell markers were quantified with qRT-PCR from punch biopsy skin samples using primers specific for mast cells (*mcpt4*), eosinophils (*embp*), neutrophils (*mpo*), basophils (*mcpt8*), T lymphocytes (*cd3*), and macrophages (*iba1*). This approach revealed that, for male mice, expression of all leukocyte markers was increased in infested skin compared to naïve skin, regardless of infestation number. Between one and four infestations, only markers for eosinophils (Fig 3F), neutrophils (Fig 3I), basophils (Fig 4C), and T lymphocytes (Fig 4F) were increased in males. In female mice, markers for eosinophils (Fig 3F), neutrophils (Fig 3I), basophils (Fig 4C), T lymphocytes (Fig 4F), and macrophages (Fig 4I) increased in mice infested four times when compared to naïve samples; however, statistically significant increases were not detected for any leukocyte markers between one and four exposures. Mast cell and macrophage markers were not increased between one and four infestations in either sex. The mast cell quantification matched our observations with toluidine blue staining (Fig 3A-B), that a robust mast cell infiltrate was not observed by staining. Similarly, for macrophages, immunohistochemistry against IBA1 did not show an increased macrophage influx to the larval bite site (Fig 4G-H; Tables 1-2). Taken together, these findings suggest that basophils, eosinophils, neutrophils, and T lymphocytes may be important contributors in driving ATR against *I. scapularis* larvae.

### Larvae fed on tick-sensitized hosts ingest greater numbers of leukocytes

Given the significant reduction in larval feeding success on sensitized *P. leucopus*, we next asked if tick resistance led to appreciable histologic changes within the midgut of feeding larvae. Larvae were collected from mice that were naïve or that had been serially infested for histologic examination. Hematoxylin and eosin staining revealed that the quantity of nucleated cells within the midgut of replete larvae increases significantly with each infestation previously experienced by the host (Fig 5; Tables 1-2). Average nucleated cells within a 150 x 150 µm area of the midgut increased from 1.2 ± 0.2 to 18.5 ± 1.7 for larvae fed on males and from 3.1 ± 0.4 to 15.6 ± 1.7 for larvae fed on females between first and fourth infestations (Fig 5G-H). The majority of the nucleated cells were characterized by segmented nuclei (polymorphonuclear) and are morphologically distinct from the mononuclear midgut epithelial cells and hemocytes of the tick (38–40), indicating that they are of host origin. We were unable to detect host cell populations by immunohistochemistry or qRT-PCR, possibly due to degradation of the blood meal during digestion. Our findings demonstrate that tick resistance is associated with a shift in bloodmeal contents characterized by increased numbers of ingested host leukocytes.

**Figure 5.**
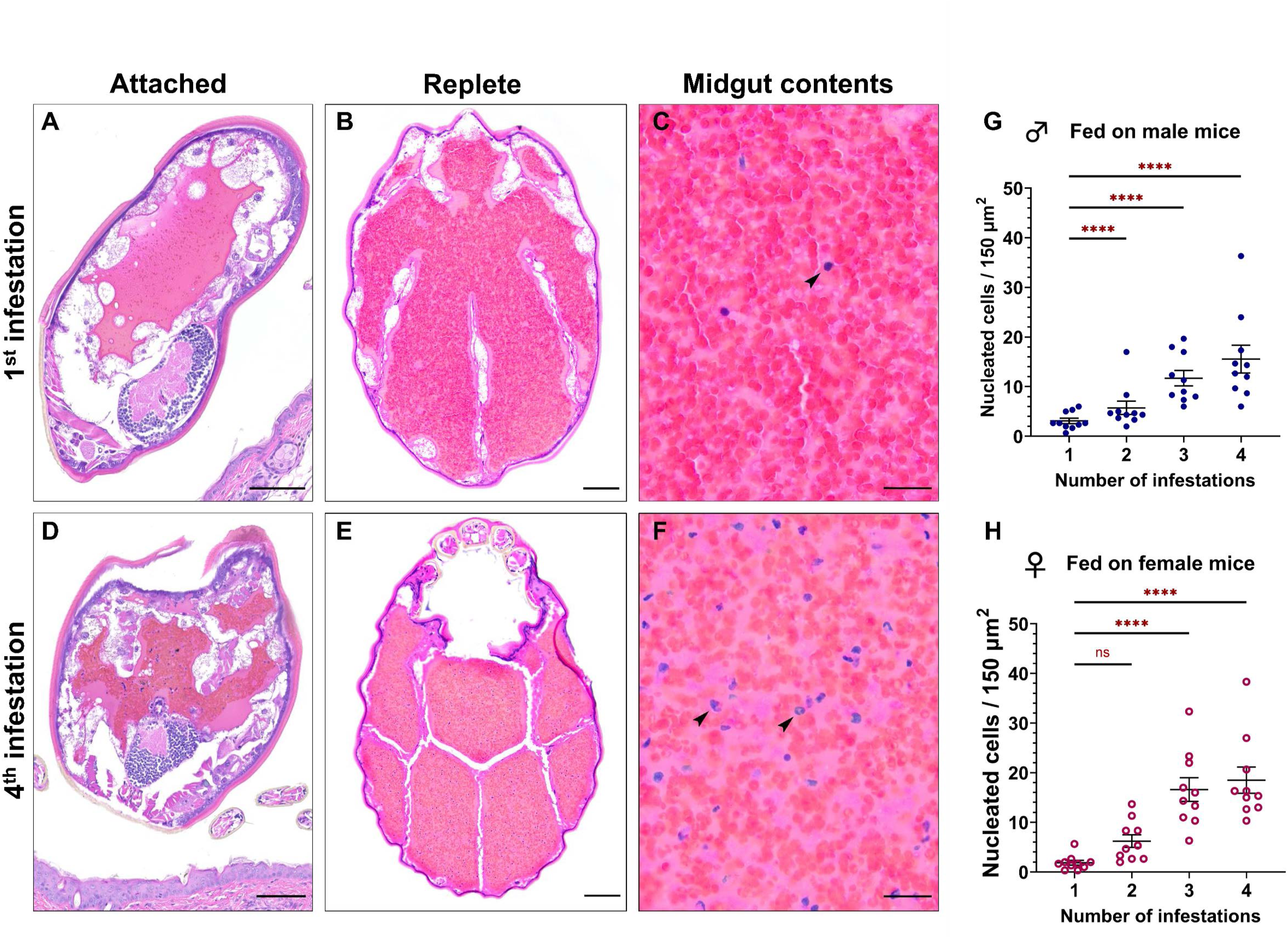
Larvae fed on tick-sensitized hosts ingest more host leukocytes. Histologic images of attached and replete larvae. (A-C) Histology of larval cross sections from ticks fed on a naïve mouse (1^st^ infestation) at 3-days post-attachment (A) and after repletion (B). (C) Nucleated cells shown in the midgut (arrow) at high magnification. (D-F) Histology of larval cross sections from ticks fed on a sensitized mouse (4^th^ infestation) at 3-days post-attachment (D) and after repletion (E). (F) Nucleated cells with multilobulated or fragmented nuclei are shown in the midgut (arrows) at high magnification. H&E stain. A-B, D-E bar = 100 μm; C, F bar = 20 μm. (G-H) Nucleated cell counts within the midguts of replete larvae fed on male (G) or female (H) mice. Each data point represents the mean nuclear count from three sites in the midgut of a single replete larva. Nucleated cell counts were analyzed using a negative binomial generalized linear mixed effect model. **p < 0.01; ****p < 0.0001. ns = not significant.

### ATR enhances larval acquisition of A. phagocytophilum and B. burgdorferi

*A. phagocytophilum* is an obligate intracellular pathogen that infects and propagates in granulocytes, particularly neutrophils, during mammalian infection (41). In contrast, neutrophils can kill *B. burgdorferi* through phagocytosis, oxidative bursts, hydrolytic enzymes, and neutrophil extracellular traps (42, 43). Since we observed a significant neutrophil component as part of the severe inflammation at the bite site on sensitized mice (Fig 3-4) and significant changes in blood meal composition within replete larvae (Fig 5), we asked if these changes could change pathogen acquisition by *I. scapularis.* To address this, we infected tick-naïve and tick-sensitized mice with either *A. phagocytophilum* or *B. burgdorferi* by needle inoculation. This approach was chosen to control for variables that are associated with tick bite transmission, including undefined infectious dose and the potential for unsuccessful tick attachment owing to mouse grooming behavior (44, 45). Naïve larvae were allowed to feed to repletion on infected mice and pathogen burdens were quantified. We found that larvae fed on previously sensitized female *P. leucopus* ingested 3.66 ± 0.90-fold more *A. phagocytophilum* than those fed on tick-naïve mice. Ticks that fed on male mice showed a similar trend (Fig 6A-B), however this difference was not statistically significant. When mice were infected with *B. burgdorferi,* the opposite trend was observed. We found that larvae fed on previously sensitized male mice ingested 46.93 ± 19.80-fold more *B. burgdorferi* than those fed on tick-naïve mice. While no difference was observed in larvae fed on naïve or sensitized female mice, there was a similar but non-significant increasing trend (Fig 6C-D; Tables 1 and 2). Taken together, this data demonstrates that ATR impacts how many pathogens are ingested by naïve *I. scapularis* larvae from infected *P. leucopus*. Notably, although sex did not impact larval feeding success, it did influence pathogen load in the tick, suggesting that undefined variables between male and female mice exhibiting ATR influence *A. phagocytophilum* and *B. burgdorferi* transmission dynamics.

**Figure 6.**
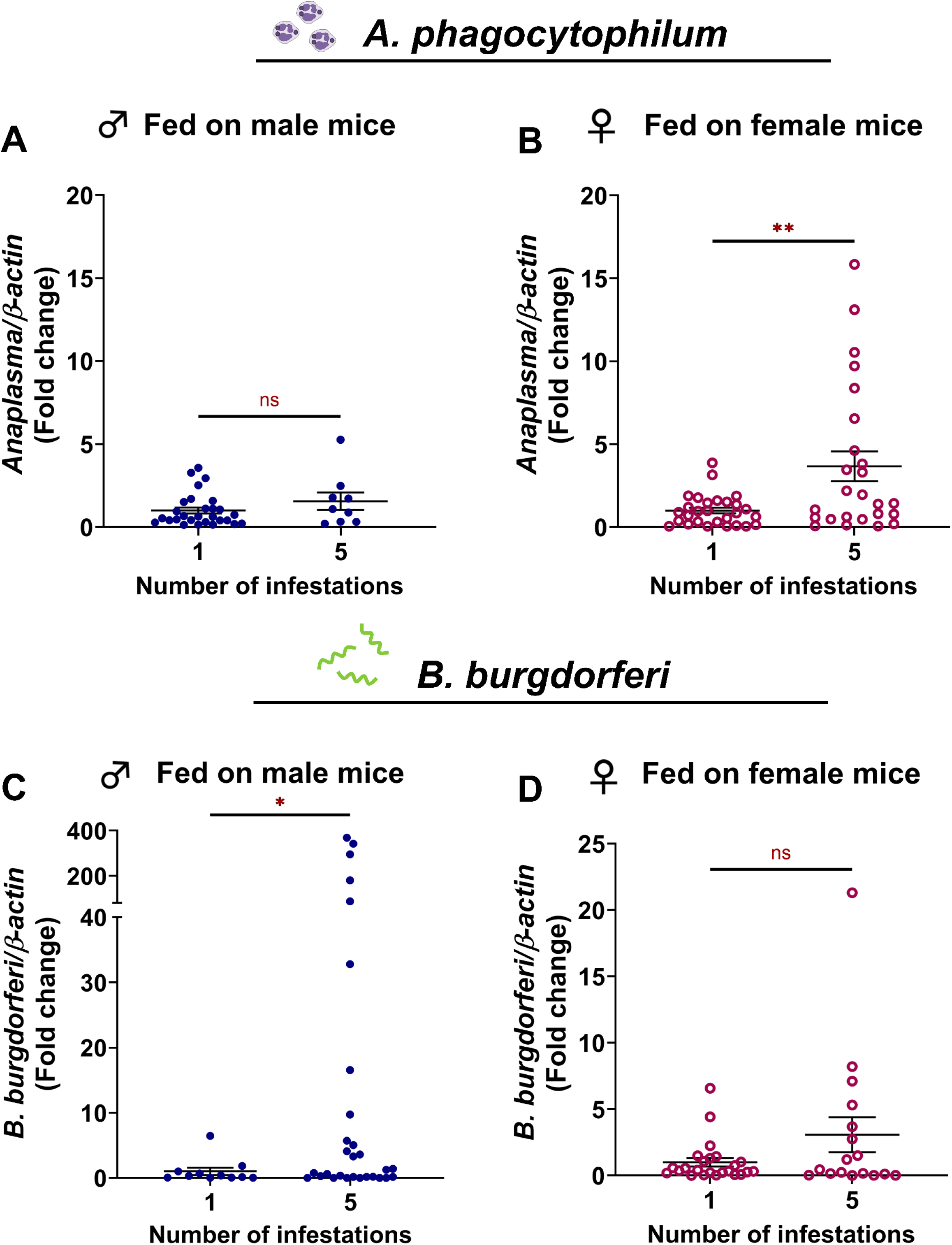
P. leucopus ATR against I. scapularis larvae alters A. phagocytophilum and B. *burgdorferi* acquisition by ticks. Naïve larvae were fed to repletion on *A. phagocytophilum*-infected (A-B) or *B. burgdorferi*-infected (C-D) male and female *P. leucopus* that were either tick-naïve or sensitized with four previous larval infestations. Pathogen burdens were assessed via qRT-PCR and normalized to tick-naïve conditions. Pathogen burdens were analyzed using a parametric Welch’s t test. Each data point represents a single replete larva. *p < 0.05; **p < 0.01. ns = not significant.

## DISCUSSION

How a host species interacts with hematophagous arthropods impacts vector competence and pathogen movement within natural systems (46), both in ticks (47–49) and in other vectors like mosquitoes (50–53). In this study, we demonstrate repeated exposure to *I. scapularis* larvae causes an increasingly severe inflammatory response in *P. leucopus*. This correlates with decreased larval feeding success characterized by fewer replete ticks and lower replete weights, which is consistent with previous reports (27, 29). We found that larvae fed on sensitized mice showed altered blood meal compositions with an increased amount ingested host leukocytes. Moreover, ATR caused increased *B. burgdorferi* and *A. phagocytophilum* ingestion by *I. scapularis* larvae in a sex-dependent manner. To our knowledge, this is the first time that ATR has been implicated in pathogen movement between native reservoir hosts and *I. scapularis* larvae.

Tick resistance in reservoir hosts has been largely overlooked in favor of using model host organisms (5, 12, 34). For example, guinea pigs exhibit an exaggerated resistance phenotype with over 80% fewer ticks feeding to repletion (20, 25, 26). For studies aiming to develop tick-targeted vaccines, near to complete cessation of tick feeding on sensitized hosts is desirable and in these cases such host organisms are appropriate models. Natural tick-host pairings do not generally elicit such a high degree of resistance (35, 49). Nevertheless, the 58% to 70% reduction in larval feeding success observed in our study when larvae are fed on tick-sensitized hosts would result in fewer ticks within the environment. Tick-borne pathogen transmission within natural systems is a density dependent phenomenon with greater numbers of infected ticks in the environment being associated with a greater risk of infection (54, 55). Previous studies have correlated reductions in questing nymphs ranging from 68% to 76% following acaracide treatment of reservoir hosts with a 53% to 96% reduction in Lyme disease risk (56–61). It’s possible that the degree of larval resistance in reservoir hosts is sufficient to influence pathogen transmission risk. This may be especially true during periods when many tick-naïve hosts are entering the population, such as the bimodal peaks of *P. leucopus* reproductive activity in spring and autumn (62). Additionally, reduced replete weight is correlated with reduced fecundity (49, 63) and post-molt size (49, 64), notable because smaller ticks are more susceptible to desiccation because of their higher surface to volume ratio (65, 66). Though these factors were not directly evaluated in our study, they may also influence the number of questing ticks within the environment. Interestingly, the reduction in larval *I. scapularis* feeding success on tick-sensitized *P. leucopus* is similar to that seen in *Dermacentor andersoni* larvae fed on deer mice (*Peromyscus maniculatus*) and cottontail rabbits (*Sylvilagus nuttallii*) (49), in *Ixodes trianguliceps* fed on bank voles (*Clethrionomys glareolus*) (67), and on larval *I. scapularis* fed on meadow voles (*Microtus pennsylvanicus*) (27). This suggests that the phenomenon of tick resistance in native hosts spans across species within natural systems and may have an underappreciated role in the natural tick-pathogen-host dynamics.

Previous work shows that *I. scapularis* nymphs elicit increasingly severe inflammation at the bite site in tick-sensitized *P. leucopus* (30). However, this does not correlate with a drop in feeding success, in contrast to what we observed with larvae. Nymphal infestations did not show a basophil influx at bite site until four days post-attachment and the majority were confined to the blood vessel lumens (30). We found basophil infiltrates at larval bite sites three days post infestation that were centered around the embedded hypostome, suggesting that basophils may play an essential role in mediating ATR against *I. scapularis* larvae. This is in agreement with previous findings in *Mus musculus* subjected to repeated infestation with *Haemaphysalis longicornis* larvae in which ablation of basophils resulted in an inability to develop tick resistance (68). This may suggest that older life stages are better able to suppress basophil infiltrate than immature tick life stages, facilitating their feeding success despite inflammation at the bite site.

Tick saliva plays an important immunosuppressive role during tick feeding (69–80). Qualitative and quantitative differences in saliva may be responsible for the differential impact of ATR on feeding success between tick life stages. Comparisons of the sialome between nymphal and adult *H. longicornis* revealed 30 proteins produced exclusively in nymph saliva, including the protease inhibitors serpin and cystatin and immunomodulatory alpha-1-acid glycoprotein 1. Seventy-four proteins were exclusively found in adult saliva, including the antioxidant catalase and immunomodulatory galectin-1 (81). Additionally, the volume of saliva produced by a feeding tick correlates with the volume of both the hemolymph and blood meal (82, 83). It is possible that the smaller volume and altered salivary content produced by *I. scapularis* larvae is not sufficient to abrogate the negative effects of the *P. leucopus* immune response, including infiltration and histamine release by basophils. Histamine and serotonin produced in sensitized hosts in response to tick attachment decrease both saliva production and feeding success in adult *Dermacentor andersoni* (84). The negative effects of the host immune response may be exacerbated in larvae due to their smaller size, higher surface to volume ratio, and limited lipid stores (65, 66).

Histamine-induced epidermal hyperplasia can prevent some ectoparasites from accessing host blood, as in the case of the northern fowl mite (*Ornithonyssus sylviarum*). Increasing inflammation and epidermal thickening separate the mite from the blood vessels of its avian host (85). It is possible that the lower feeding success we observed with larval *I. scapularis* on sensitized mice could be attributable to epidermal hyperplasia impeding access to the feeding lesion (5). The average length of larval *I. scapularis* hypostomes are 0.111 mm ± 0.037 mm (86). In our study, tick-sensitized mice exhibited sufficient epidermal thickening to potentially impact the ability of larvae to reach the feeding lesion (Tables 1-3). However, we observed that the epidermis immediately adjacent to the hypostome was often eroded or ulcerated with no appreciable displacement of the tick from the feeding lesion. It therefore remains unclear what effect, if any, epidermal hyperplasia has on larval attachment. The smaller size of the larval hypostome and cement cone may make dermal anchoring inherently less stable when compared to nymphs. However, given the epidermal loss and extension of granulocytes into the cement cone observed in this study, destabilization of the dermal-cement adhesion may be a more feasible explanation for failure to reach repletion.

The majority of studies evaluating host resistance to hematophagous arthropod feeding have focused on how inflammation impacts transmission of vector-borne pathogens from the arthropod to the host. For example, host resistance alters transmission outcomes for mosquitos-borne arboviruses (87–89), sandfly-borne leishmaniasis (90–93), and tick-transmitted pathogens (45, 94–99). Tick resistance greatly reduces host susceptibility to pathogen transmission by infected ticks, including *B. burgdorferi* (guinea pigs and non-human primates) (45, 94, 95), *Francisella tularensis* (rabbits) (98), *Babesia* spp. (cattle) (96, 97), *Anaplasma marginale* (cattle) (97), and tick-borne encephalitis virus (*M. musculus* infested with *R. appendiculatus*) (99). Even in the absence of tick resistance, inflammation at nymphal *I. scapularis* bite sites of repeatedly infested *P. leucopus* reduced rates of *B. burgdorferi* transmission from the tick to the host by 83.3% (44). To our knowledge, our study is the first to evaluate the interplay between *P. leucopus* resistance to larval *I. scapularis* and pathogen ingestion by ticks.

We observed sex-specific variations in both tick resistance and pathogen transmission from mice to *I. scapularis*. In general, fewer larvae fed to repletion on females than males, consistent with previous reports (27, 100–103). We also found that larvae fed on previously sensitized, female *P. leucopus* ingested more *A. phagocytophilum* than those fed on tick-naïve mice. Enhancement of *A. phagocytophilum* acquisition from tick-sensitized female mice is consistent with the histologic changes that characterize ATR in both the host skin and replete ticks. *A. phagocytophilum* infects and replicates within neutrophils, eosinophils, and monocytes (104). As such, the increase in granulocytes at both the host bite site and within the replete larval midgut would be expected to enhance larval ingestion of *A. phagocytophilum* from tick-sensitized hosts. However, similar enhancement of *A. phagocytophilum* transmission was not observed in male mice, suggesting that additional factors must be at play.

Sex differences in immunity, pathogen kinetics, and host-pathogen interactions may also influence pathogen acquisition from infected hosts. General immunity differences between the sexes has been previously reported with both female mice and humans mounting more robust innate and adaptive immune responses than males (105–108). As a result, females tend to be less susceptible to parasitic and viral infections (109–111). Differences in susceptibility also extend to tick-borne pathogens. Male C57BL/6 mice exhibit increased susceptibility to *A. phagocytophilum* infection with up to a 1.85-fold increase in infected neutrophils and significantly greater splenomegaly compared to infected females (112). If *A. phagocytophilum* burdens were sufficiently high in male *P. leucopus*, it is possible that any boost in acquisition provided by tick resistance could be muted.

Another possibility not explored in this study is the influence of infection on the inflammatory infiltrate at larval attachment sites. *A. phagocytophilum* can modify the activity of neutrophils by suppressing respiratory bursts (113), dysregulating degranulation (114), and interfering with surface selectin expression resulting in impaired transmigration from the vasculature (115, 116). Given its ability to modulate granulocyte behavior, it is possible that *A. phagocytophilum* could reduce leukocyte transmigration to larval attachment sites. Changes in the inflammatory infiltrate could help explain why tick-sensitized male mice did not show enhanced transmission of *A. phagocytophilum* to larvae, especially if higher *A. phagocytophilum* burdens in males leads to more profound alterations in neutrophil behavior. However, additional studies would be necessary to evaluate whether infection can influence the inflammatory response to ticks.

Interestingly, the sex-specific effects on pathogen transmission were reversed in the case of *B. burgdorferi*. We found that larvae fed on previously sensitized male mice acquired more bacteria than those fed on naïve *P. leucopus*, and no difference was observed between larvae fed on naïve or tick-sensitized females. This finding was unexpected given previous studies demonstrating that prior tick exposure reduces the susceptibility of *P. leucopus* to *B. burgdorferi* transmission by infected nymphs (44). One possible explanation is that inflammatory mediators and/or DAMPs, such as reactive oxygen species (ROS) and L-serine in serum, may act as chemoattractants for *B. burgdorferi*, as is seen with *H. pylori* (117) and other enteric pathogens including *S. enterica*, *E. coli*, and *C. koseri* (118, 119). Sex differences in the production of inflammatory mediators could conceivably lead to differences in chemotactic responses. For instance, males tend to produce more histamine in response to cutaneous allergens (120), have higher basal levels of ROS, and less efficient antioxidant mechanisms (121–123). Alternatively, it is possible that the tendency toward a less robust immune response in males could allow for more effective transmission of *B. burgdorferi* to feeding larvae. In *M. musculus* infected with *B. burgdorferi*, males showed both a higher percentage of infected tissues and higher cutaneous and visceral spirochete burdens (124). Pathogen transmission and acquisition dynamics may be further complicated by the immunomodulatory effects exerted by *B. burgdorferi* as it attempts to subvert the host immune response. *B. burgdorferi* suppresses host IgG responses to facilitate its dissemination and persistence (125). Consequentially, the IgG response to unrelated antigens, such as the SARS-CoV-2 spike protein, is also impaired (126). If *B. burgdorferi* can immunomodulate *P. leucopus* in a similar manner, it could alter the host response to tick attachment and lead to unexpected tick-pathogen interactions.

The observation that ATR enhances larval *B. burgdorferi* acquisition from male *P. leucopus* contrasts with previous findings in *M. musculus* in which prior tick sensitization of *B. burgdorferi*-infected C57BL/6J mice was correlated with decreased pathogen ingestion by *I. scapularis* larvae (127). Our findings suggest that tick-host-pathogen interactions observed in *M. musculus* cannot necessarily be translated to native host species like *P. leucopus*. Why transmission differences exist between *M. musculus* and *P. leucopus* is unclear. Both hosts have unique genetic backgrounds (inbred C57BL/6 mice vs. outbred *P. leucopus*) which in turn influence the immune response. For instance, T cells in C57/BL6 mice favor production of Th1 cytokines such as IFN-γ (128). Differences in immunity may alter the tick-host-pathogen relationship with implications for pathogen acquisition. Alternatively, some tick salivary proteins act as chemoattractants for *B. burgdorferi* (129, 130) and *I. scapularis* alters its sialome composition depending on the host species it feeds upon (131). Differences in mouse species and tick-sensitization status likewise may trigger variation in the sialome of feeding larvae, and this in turn may influence the presence and/or abundance of chemoattracts. Although it is clear that ATR, host species, and sex-specific differences influence pathogen ingestion by ticks, additional studies are needed to determine whether this can also influence pathogen maintenance through the molt and/or transmission to naïve hosts. Additionally, while tick-sensitization enhances initial pathogen ingestion by larvae, the negative impact of ATR on larval feeding success would result in fewer infected *B. burgdorferi* infected ticks overall. This would be expected to reduce pathogen transmission risk in natural systems.

One limitation of this study is the route in which naïve or sensitized mice were infected. We chose needle inoculation to minimize uncontrolled variables that are associated with infection by tick bite, including undefined pathogen dose and the potential for unsuccessful tick attachment. With this approach, we were able to study how ATR responses affected pathogen movement into feeding larvae in a controlled, experimental setting. We acknowledge that a naïve mouse would not exist in the wild, as the only route for infection is through tick bite. However, this type of comparison is useful for isolating and examining immunological variables impacting pathogen ingestion by ticks.

Our findings showcase the complexity of factors influencing ATR and highlight that there is still much to be discovered. Sex, host species, degree of prior tick sensitization, and tick life stage all influence the presence and severity of ATR with implications for the vector-pathogen life cycles. Additionally, our study highlights the importance of including both sexes in ATR and tick-borne pathogen studies. Future studies accounting for the various factors influencing tick resistance and pathogen movement in natural systems will provide greater insight into the mechanistic underpinnings of ATR.

## MATERIALS AND METHODS

### Animal models

All mouse experiments were conducted in accordance with the guidelines and protocols approved by the American Association for Accreditation of Laboratory Animal Care (AAALAC) and by the Office of Campus Veterinarian at Washington State University (Animal Welfare Assurance A3485-01, IACUC-approved protocol #6097). White-footed mice (*P. leucopus*) were originally obtained from the Peromyscus Genetic Stock Center (University of South Carolina) to start a laboratory colony maintained at Washington State University. Mice were maintained in an AAALAC-accredited facility at Washington State University in Pullman, WA. All procedures were approved by the Washington State University Biosafety and Animal Care and Use Committees.

Pathogen-free *I. scapularis* larvae were obtained from Oklahoma State University (Stillwater, OK, USA). Ticks were kept in glass vials in an incubator maintained at 23°C and 95– 100% relative humidity with 16:8-h light:dark photoperiods.

### Infestation Procedure

Eight tick-naïve mice of each sex between 6 and 12 weeks of age were infested with *I. scapularis* larvae four successive times separated by 2-week tick-free intervals after the primary and tertiary infestation and 1-week tick-free intervals after the secondary infestation. For each infestation, mice were anesthetized (1-2% isoflurane, 1.0 liter per minute oxygen flow rate) and 100 larval ticks were placed around and between the ears and over the shoulders. Mice were maintained under anesthesia for 20 minutes following tick placement to allow time for attachment. Following recovery from anesthesia, mice were housed individually in specialized multi-cage systems to prevent tick escape. Systems included an innermost cage containing a raised metal rack suspended above 1 to 2 cm of water and a larger outer cage containing approximately 2 cm of water. Replete ticks were collected daily following natural detachment over the course of 7 days. Ticks were either weighed and maintained to monitor post-feeding survival and molting or were saved in 10% neutral buffered formalin for histologic examination. A subset of male and female mice were euthanized on day 3 post-tick placement during the primary and quaternary infestations to examine leukocyte infiltrates at tick bite sites. For leukocyte quantification by qRT-PCR, four to six tick attachment sites per group were collected from the pinnae using a 4mm sterile dermal biopsy punch (McKesson, 16-9840), placed in RNAlater Stabilization Solution (Thermo Fisher Scientific, AM7020), and frozen at −80°C. For comparison, biopsies from equivalent sites on the pinna were collected from control mice which were never infested with ticks. Bite sites collected from the head and ears for histologic examination were placed in 10% neutral buffered formalin for 24 hours prior to processing.

### Bacterial maintenance and inoculation

*A. phagocytophilum* strain HZ was cultured in HL60 cells (ATCC, CCL-240) maintained in Roswell Park Memorial Institute (RPMI) 1640 medium supplemented with 1x Glutamax (Gibco, 35050061) and 10% heat-inactivated fetal bovine serum (FBS; Atlanta Biologicals, S11550). Cultures were maintained at a concentration between 1 x 10^5^ and 1 x 10^6^ cells per ml at 37°C and 5% CO_2_. On the day of infection, *A. phagocytophilum* was quantified using the previously described method (132, 133) and was liberated from HL60 host cells via syringe lysis with a 27-gauge needle. Age-matched male and female *P. leucopus* mice were infected intraperitoneally with 1 x 10^7^ *A. phagocytophilum* HZ in 100 µL of PBS (Intermountain Life Sciences, BSS-PBS). Mice were rested for three days to allow them to reach peak bacteremia (134). On day three post-infection, between 25 and 50 µL of infected blood was collected from the lateral saphenous vein of each mouse, and *A. phagocytophilum* burdens were assessed via quantitative PCR (16S relative to mouse β-actin) to confirm infection and for the purpose of burden-matching (132, 135, 136). The following day, infected mice were infested with 100 larval ticks as described above. For analysis, ticks fed during quaternary infestations were normalized to ticks fed on burden-matched hosts during primary infestations.

*B. burgdorferi* B31 (strain MSK5) was cultured in modified Barbour-Stoenner-Kelly II (BSK-II) medium supplemented with 6% normal rabbit serum (NRS; Pel-Freez; 31126-5) and maintained at 37°C and 5% CO_2_. Spirochete growth was monitored daily using dark-field microscopy. Prior to infection, PCR was used to screen the plasmid profiles of *B. burgdorferi* cultures (137). Male and female age-matched *P. leucopus* mice were infected intradermally with 1 x 10^5^ low-passage *B. burgdorferi* B31 MSK5 in 100 µl 50:50 PBS/NRS. Mice were rested for four weeks to allow spirochetes time to disseminate (138), then infested with 100 larval ticks as described above.

### Histology and Immunohistochemistry

Skin samples and whole replete larval ticks were fixed in 10% neutral buffered formalin for 24 hours and embedded in paraffin. 5 μm sections were collected and either stained with H&E for routine histopathology, Toluidine blue for detection of mast cells, Luna stain for detection of eosinophils, or were deparaffinized using clearing agent Clear-Rite 3 (Medix Corp, 6901) and rehydrated for immunohistochemistry. Heat-induced epitope retrieval (HIER) was performed by heating slides in a microwave at 700 watts for 20 minutes in either citrate buffer, pH 6.0 (Abcam, ab64214) or EDTA disodium salt dihydrate solution adjusted to pH 8.0 (Invitrogen, 15576028). Following a 30-minute room-temperature incubation with high protein blocking buffer (ThermoFisher, 00-4952-54), samples were incubated overnight at 4°C with one of the following antibodies diluted to 1:100 in high protein blocking buffer: α-myeloperoxidase polyclonal antibody (ThermoFisher, PA5-16672) for neutrophils, IBA1 polyclonal antibody (Fisher, PIPA527436) for macrophages, or MCP-8 monoclonal antibody (BioLegend, 647401-BL) for basophils. Following room-temperature 10-minute 3% hydrogen peroxide block, samples were incubated with one of two secondary antibodies at room temperature for 1 hour: goat pAB to rabbit IgG conjugated to HRP (Abcam, ab97051) at 1:200 dilution, or goat pAB to rat IgG conjugated to HRP (ThermoFisher, 31470) at 1:250 dilution. Slides were treated with the AEC Substrate Kit (Abcam, ab64252) with extended incubation for 25 minutes. Slides were stained with Mayer′s Hematoxylin Solution (H&E, Sigma, MHS32-1L), and cover-slipped using aqueous mounting medium for IHC (Abcam, ab64230).

For detection of macrophages, the above protocol was slightly modified. Peroxide block was performed prior to antigen retrieval using 0.5% hydrogen peroxide in 85% ethanol incubated at room temperature for 30 minutes. For heat-induced epitope retrieval, the microwave was set to 800 watts.

For detection of T lymphocytes, deparaffinization and immunohistochemistry were performed by the Washington Animal Disease Diagnostic Laboratory Histology Core using the Ventana Discovery Ultra automated stainer (Roche). Heat-induced epitope retrieval (HIER) was performed by heating slides to 95°C in ULTRA CC1 antigen retrieval solution (Roche, 950124) for 56 minutes. Slides were incubated for 24 minutes at room temperature with CD3 polyclonal antibody (Dako, A0452) diluted to 1:400 in Antibody Dilution Buffer (Roche, ADB250). To visualize immunoreactivity, slides were treated using the UltraView Universal Alkaline Phosphatase Red Detection Kit (Roche, 760-501) per manufacturer instructions.

### Histologic Evaluation of Larval Attachment Sites

Histologic evaluation of bite sites was conducted with the aid of a board-certified veterinary anatomic pathologist. Severity was defined as such: Mild – small numbers of inflammatory cells are present, and inflammation is confined to the dermis; Moderate – moderate numbers of inflammatory cells are present and may extend into the superficial subcutis and/or skeletal muscle; Severe – numerous inflammatory cells are present with extension deep into the subcutis and skeletal muscle and/or crossing of the auricular cartilage. To assess epidermal hyperplasia, measurements of epidermal thickness were collected adjacent to sites of larval attachment using an ocular micrometer at 400x magnification. Measurements were taken in duplicate from each site and averaged. As a control, epidermal measurements were collected from the same animal at the same anatomical location away from sites of tick attachment and from uninfested control animals at comparable locations. Epidermal measurements were taken perpendicular to and beginning at the basement membrane.

### Analysis of Larval Midgut Contents

To quantify nucleated cells within replete larval midguts, ten replete larvae from each infestation on both male and female hosts were evaluated. Three 150 x 150 µm images were collected per larva centered within a separate midgut lobule. Nuclei were quantified in each image using ImageJ (parameters: pixil count (50-infinity); circularity 0.20-100).

### Quantitative Reverse Transcriptase Polymerase Chain Reaction (qRT-PCR)

RNA was extracted from skin biopsy samples and from replete ticks via the Direct-zol RNA MicroPrep kit (Zymo Research, R2062). cDNA was synthesized from 300 to 500 ng total RNA was performed using the Verso cDNA Synthesis Kit (ThermoFisher, AB-1453). Bacterial burden and leukocyte transcripts were assessed by qRT-PCR with iTaq universal SYBR green Supermix (Bio-Rad; 1725125) using the primers listed in Table 4 and cycle conditions as recommended by the manufacturer. For leukocyte marker quantification, values were converted to log base 10.

### Statistical analysis

Statistical tests were done using the program R (version 4.2.2). Data was evaluated independently for each host sex. Number of larvae recovered from naïve and sensitized hosts and larval midgut leukocyte counts were analyzed using negative binomial generalized linear mixed effect models with the function *glmer.nb* (package *MASS*). Models treated the host’s prior tick exposure as a fixed effect and host ID, infestation time point, and tick cohort as random effects. The proportion of larvae molting successfully was analyzed using a generalized linear mixed-effects model with the function *glmer* (package *MASS*). The time taken to molt was analyzed using a negative binomial generalized linear mixed effect model with the function *glmer.nb* (package *MASS*). Models treated the host’s prior tick exposure as fixed effects, and host ID, host sex, and tick cohort as random effects. Replete larval weights was analyzed using a linear mixed effects regression model with the function *lmer* (package *lmerTest*), treating the host’s prior tick exposure as a fixed effect and host ID, infestation time point, and tick cohort as random effects. Epidermal thickness at larval attachment sites was analyzed using a linear mixed effects regression model with the function *lmer* (package *lmerTest*), treating the host’s prior tick exposure as a fixed effect and biopsy sample ID as a random effect. Leukocyte marker quantification at larval bite sites by qRT-PCR was analyzed using a linear regression model with the function *lm* (included in base R). Post-hoc comparisons were performed using the function *emmeans* (package *emmeans*) for larval replete weights, molting success, molting time, midgut leukocyte counts, and qRT-PCR quantification of leukocyte markers. *A. phagocytophilum* and *B. burgdorferi* burdens were analyzed using a parametric Welch’s t test. For all analyses, the cutoff for statistical significance was set at a P value of <0.05. Descriptive statistics for each experiment are summarized in Table 1. Statistical models and results are summarized in Table 2.

## ACKNOWLEDGEMENTS

We are grateful to Jon Skare (Texas A&M Health Science Center) for providing *Borrelia burgdorferi* B31 clone MSK5, the Washington Animal Disease Diagnostic Laboratory Histology Core and Susan Noh for their assistance with developing and refining immunohistochemistry protocols, and Oklahoma State University and Biodefense and Emerging Infectious Diseases Resources for providing *Ixodes scapularis* larvae.

## AUTHOR CONTRIBUTIONS

E.A.F., J.P.O, and D.K.S. designed the study. E.A.F., C.J.L., K.L.R., and E.R-Z. performed the experiments. E.A.F., J.R.A, J.P.O., and D.K.S. analyzed the data. All authors provided intellectual input into the study. E.A.F and D.K.S. wrote the manuscript. All authors contributed to editing.

## FIGURE LEGENDS

**Supplemental Figure 1. High magnification images of larval attachment sites on tick-naïve and tick-sensitized mice**. Histological examination of male and female mouse biopsies from (A-B) primary infestations, and (C-D) quaternary infestations. H&E stain. Bar = 50 μm.

