## Supplemental Figure 1 for "Acquired tick resistance in *Peromyscus leucopus* alters *Ixodes scapularis* infection"

**Supplemental Figure 1. High magnification images of larval attachment sites on tick-naive and tick-sensitized mice**

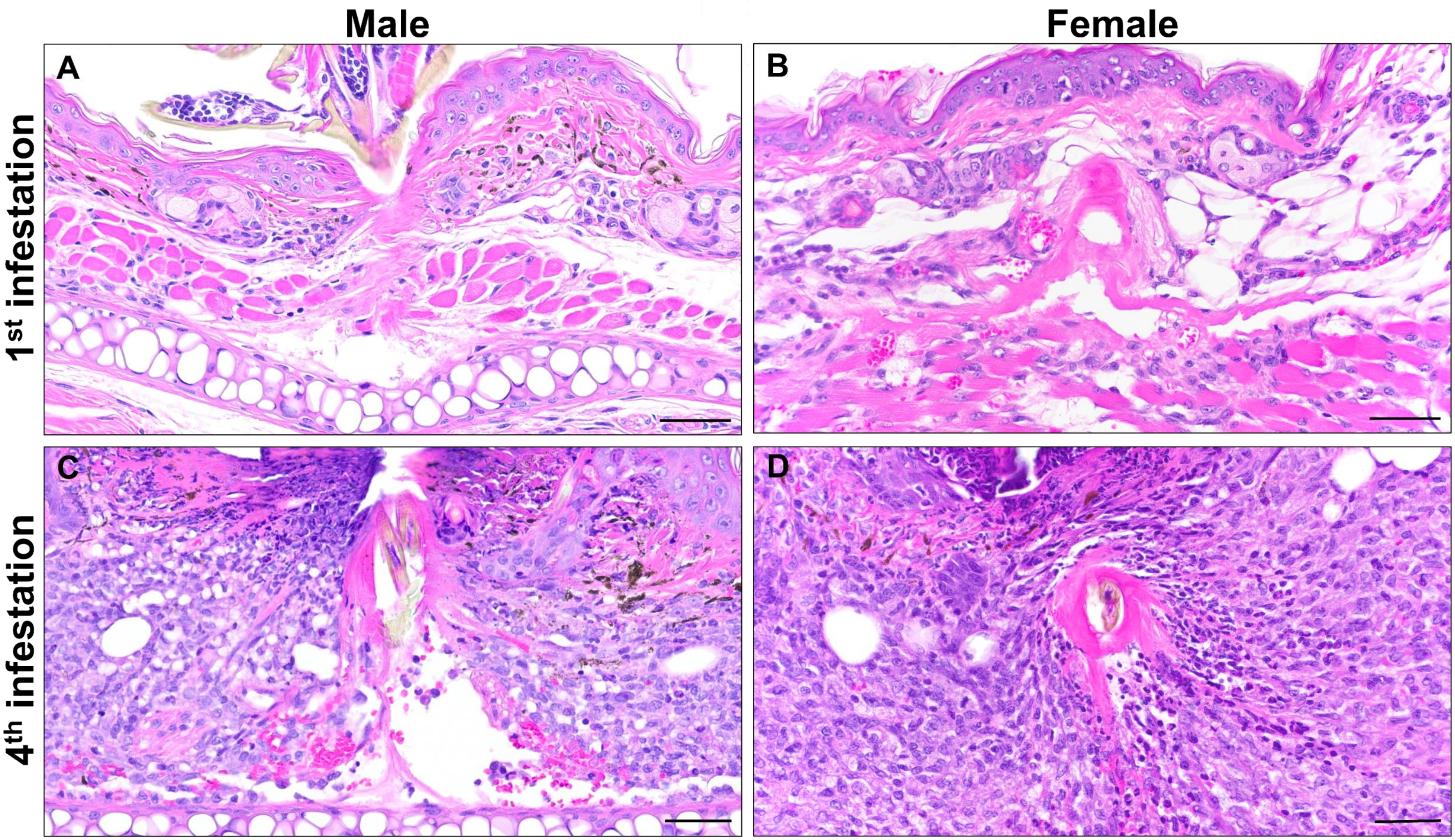
