## Supplementary material for "Acquired tick resistance in *Peromyscus leucopus* alters *Ixodes scapularis* infection": Table 1

**Table 1. Descriptive statistics for experiments in this study**

| Feeding Success | | | | |
|---|---|---|---|---|
| Group | Host sex | n (mice) | Mean (larvae reaching repletion) | Standard error |
| 1° | Male | 8 | 19.125 | 1.9220292 |
| 2° | Male | 8 | 8.5 | 1.2100767 |
| 3° | Male | 8 | 7.375 | 0.8647357 |
| 4° | Male | 8 | 5.75 | 1.0307764 |
| 1° | Female | 8 | 12.75 | 1.5089022 |
| 2° | Female | 8 | 5.625 | 0.7055267 |
| 3° | Female | 8 | 6.25 | 0.9013878 |
| 4° | Female | 8 | 5.375 | 1.0511473 |

| Larval Replete Weight | | | | |
|---|---|---|---|---|
| Group | Host sex | n (larvae) | Mean (mg) | Standard error |
| 1° | Male | 58 | 0.4560345 | 0.009844341 |
| 2° | Male | 31 | 0.4254839 | 0.008673761 |
| 3° | Male | 21 | 0.4190476 | 0.009221389 |
| 4° | Male | 22 | 0.3768182 | 0.008537829 |
| 1° | Female | 72 | 0.5015278 | 0.008906706 |
| 2° | Female | 25 | 0.3852 | 0.009951884 |
| 3° | Female | 19 | 0.3952632 | 0.015320521 |
| 4° | Female | 17 | 0.3652941 | 0.015953002 |

| Larvae molting successfully | | | | |
|---|---|---|---|---|
| Group | Host sex | n (larvae) | n (molted) | n (died) |
| 1° | Male | 58 | 36 | 22 |
| 2° | Male | 32 | 31 | 1 |
| 4° | Male | 22 | 17 | 5 |
| 1° | Female | 58 | 43 | 15 |
| 2° | Female | 25 | 17 | 8 |
| 4° | Female | 17 | 14 | 3 |

| Time taken to molt | | | | |
|---|---|---|---|---|
| Group | Host sex | n (larvae) | Mean (days) | Standard error |
| 1° | Male | 36 | 31.33333 | 1.110555 |
| 2° | Male | 31 | 27.77419 | 1.068322 |
| 4° | Male | 17 | 28.52941 | 2.530865 |
| 1° | Female | 43 | 37.13953 | 1.1199299 |
| 2° | Female | 17 | 35.05882 | 1.0963529 |
| 4° | Female | 14 | 37.28571 | 0.9164466 |

| Epidermal thickness at larval attachment sites | | | | |
|---|---|---|---|---|
| Group | Host sex | n (biopsy sites) | Mean (µm) | Standard error |
| Uninfested | Male | 4 | 12.4 | 1.237509 |
| 1° | Male | 9 | 25.16667 | 1.99596 |
| 4° | Male | 3 | 57.66667 | 15.736638 |
| Uninfested | Female | 4 | 17.2 | 0.6590036 |

| | | | | |
|---|---|---|---|---|
| 1° | Female | 3 | 36.8 | 6.5057923 |
| 4° | Female | 2 | 88 | 15.2210381 |

| qRT-PCR analysis of leukocytes at tick attachment sites | | | | |
|---|---|---|---|---|
| Mast cells | | | | |
| Group | Host sex | n (biopsy sites) | Mean ($Log_{10}$ fold change) | Standard error |
| Uninfested | Male | 6 | -4.99451E-16 | 0.08545105 |
| 1° | Male | 6 | 0.4374969 | 0.08322087 |
| 4° | Male | 6 | 0.6567471 | 0.03293417 |
| Uninfested | Female | 6 | -2.52112E-16 | 0.1211938 |
| 1° | Female | 5 | 0.3574229 | 0.1603691 |
| 4° | Female | 4 | 0.4992081 | 0.2857658 |
| Eosinophils | | | | |
| Group | Host sex | n (biopsy sites) | Mean ($Log_{10}$ fold change) | Standard error |
| Uninfested | Male | 6 | 3.00684E-16 | 0.09870745 |
| 1° | Male | 6 | 1.172512 | 0.07255767 |
| 4° | Male | 6 | 1.59295 | 0.10317154 |
| Uninfested | Female | 6 | 5.41234E-16 | 0.124447 |
| 1° | Female | 5 | 0.4727174 | 0.1953576 |
| 4° | Female | 4 | 1.062887 | 0.3395693 |
| Neutrophils | | | | |
| Group | Host sex | n (biopsy sites) | Mean ($Log_{10}$ fold change) | Standard error |
| Uninfested | Male | 6 | 2.1684E-16 | 0.08332515 |
| 1° | Male | 6 | 1.329047 | 0.08833063 |
| 4° | Male | 6 | 1.818221 | 0.10354367 |
| Uninfested | Female | 6 | 1.11485E-15 | 0.1213482 |
| 1° | Female | 5 | 0.6042675 | 0.2160771 |
| 4° | Female | 4 | 1.189319 | 0.3648499 |
| Basophils | | | | |
| Group | Host sex | n (biopsy sites) | Mean ($Log_{10}$ fold change) | Standard error |
| Uninfested | Male | 6 | -4.36572E-17 | 0.074652 |
| 1° | Male | 6 | 1.024505 | 0.07917496 |
| 4° | Male | 6 | 1.497624 | 0.09525748 |
| Uninfested | Female | 6 | -5.73619E-16 | 0.08056014 |
| 1° | Female | 5 | 0.608482 | 0.16920564 |
| 4° | Female | 4 | 1.230711 | 0.34311808 |
| T Lymphocytes | | | | |
| Group | Host sex | n (biopsy sites) | Mean ($Log_{10}$ fold change) | Standard error |
| Uninfested | Male | 6 | -2.17419E-16 | 0.08767389 |
| 1° | Male | 6 | 0.7400321 | 0.06777187 |
| 4° | Male | 6 | 1.226196 | 0.11160867 |
| Uninfested | Female | 6 | -1.44329E-15 | 0.113545 |

| | | | | |
|---|---|---|---|---|
| 1° | Female | 5 | 0.5356327 | 0.1984644 |
| 4° | Female | 4 | 1.162979 | 0.3608239 |
| Macrophages | | | | |
| Group | Host sex | n (biopsy sites) | Mean (Log$_{10}$ fold change) | Standard error |
| Uninfested | Male | 6 | -1.9718E-16 | 0.0648939 |
| 1° | Male | 6 | 0.2613944 | 0.08478032 |
| 4° | Male | 6 | 0.3231055 | 0.05255204 |
| Uninfested | Female | 6 | 2.31295E-16 | 0.04307114 |
| 1° | Female | 5 | 0.8034491 | 0.15236876 |
| 4° | Female | 4 | 1.022749 | 0.3237656 |
| Nucleated cell counts within larval midguts | | | | |
| Group | Host sex | n (larvae) | Mean (nucleated cells) | Standard error |
| 1° | Male | 10 | 1.233333 | 0.2072443 |
| 2° | Male | 10 | 6.233333 | 0.7829848 |
| 3° | Male | 10 | 16.6 | 1.4436544 |
| 4° | Male | 10 | 18.5 | 1.6800041 |
| 1° | Female | 10 | 3.1 | 0.3816179 |
| 2° | Female | 10 | 5.733333 | 0.8508729 |
| 3° | Female | 10 | 11.7 | 0.9955649 |
| 4° | Female | 10 | 15.566667 | 1.6599982 |

| *A.phagocytophilum* burdens in replete larvae | | | | |
|---|---|---|---|---|
| Group | Host sex | n (larvae) | Mean (fold change) | Standard error |
| 1° | Male | 28 | 1 | 0.1851 |
| 5° | Male | 9 | 1.558 | 0.5302 |
| 1° | Female | 28 | 1 | 0.1772 |
| 5° | Female | 25 | 3.663 | 0.8956 |

| *B. burgdorferi* burdens in replete larvae | | | | |
|---|---|---|---|---|
| Group | Host sex | n (larvae) | Mean (fold change) | Standard error |
| 1° | Male | 11 | 1 | 0.5702 |
| 5° | Male | 29 | 46.93 | 19.8 |
| 1° | Female | 24 | 1 | 0.313 |
| 5° | Female | 17 | 3.075 | 1.307 |
