## Supplementary material for "Acquired tick resistance in *Peromyscus leucopus* alters *Ixodes scapularis* infection": Table 2

**Table 2. Methodology and results for statistical analyses**

| Feeding Success | | | | | |
|---|---|---|---|---|---|
| Negative binomial generalized linear mixed effect model: Proportion of larvae reaching repletion ~ infestation number + (1 \| larval cohort + infestation date + host ID) | | | | | |
| Comparisson | Host sex | β | Standard error | Z ratio | P-value |
| 2° vs 1° | Male | -0.893 | 0.199 | -4.488 | <.0001 |
| 3° vs 1° | Male | -0.9721 | 0.182 | -5.344 | <.0001 |
| 4° vs 1° | Male | -1.2136 | 0.199 | -6.097 | <.0001 |
| 3° vs 2° | Male | -0.0791 | 0.224 | -0.353 | 0.9849 |
| 4° vs 2° | Male | -0.3206 | 0.22 | -1.456 | 0.4641 |
| 4° vs 3° | Male | -0.2415 | 0.233 | -1.038 | 0.727 |
| 2° vs 1° | Female | -0.9043 | 0.194 | -4.655 | <.0001 |
| 3° vs 1° | Female | -0.8253 | 0.245 | -3.364 | 0.0043 |
| 4° vs 1° | Female | -1.0363 | 0.212 | -4.891 | <.0001 |
| 3° vs 2° | Female | 0.0791 | 0.255 | 0.31 | 0.9897 |
| 4° vs 2° | Female | -0.132 | 0.243 | -0.544 | 0.9482 |
| 4° vs 3° | Female | -0.2111 | 0.263 | -0.802 | 0.8537 |

| Larval Replete Weight | | | | | |
|---|---|---|---|---|---|
| Linear mixed effects regression model: Larval replete weight ~ infestation number + (1 \| larval cohort + infestation date + host ID) | | | | | |
| Comparisson | Host sex | β | Standard error | T-value | P-value |
| 2° vs 1° | Male | 9.31E-04 | 1.44E-02 | 0.065 | 0.948 |
| 3° vs 1° | Male | -4.25E-03 | 1.63E-02 | -0.261 | 0.795 |
| 4° vs 1° | Male | -7.19E-02 | 2.74E-02 | -2.626 | 0.027 |
| 2° vs 1° | Female | -0.11617 | 0.01546 | -7.515 | <.0001 |
| 3° vs 1° | Female | -0.1086 | 0.01975 | -5.499 | 0.00054 |
| 4° vs 1° | Female | -0.15202 | 0.02048 | -7.422 | <.0001 |

| Larvae molting successfully | | | | | |
|---|---|---|---|---|---|
| Generalized linear mixed-effects model: Molt status ~ infestation number + (1 \| host ID + host sex + experimental replicate) | | | | | |
| Comparisson | Host sex | β | Standard Error | Z ratio | P-value |
| 2° vs 1° | Male | 1.228 | 1.16 | 1.061 | 0.5386 |
| 4° vs 1° | Male | 0.472 | 0.786 | 0.6 | 0.8199 |
| 4° vs 2° | Male | -0.756 | 1.24 | -0.609 | 0.8155 |
| 2° vs 1° | Female | -0.312 | 0.561 | -0.556 | 0.8435 |
| 4° vs 1° | Female | 0.484 | 0.737 | 0.656 | 0.7888 |
| 4° vs 2° | Female | 0.796 | 0.8 | 0.995 | 0.5802 |

| Time taken to molt | | | | | |
|---|---|---|---|---|---|
| Negative binomial generalized linear mixed effect model: Time taken to molt ~ infestation number + (1 \| host ID + host sex + experimental replicate) | | | | | |
| Comparisson | Host sex | β | Standard error | Z ratio | P-value |
| 2° vs 1° | Male | -0.0497 | 0.0655 | -0.759 | 0.7282 |
| 4° vs 1° | Male | -0.0229 | 0.0757 | -0.303 | 0.9508 |
| 4° vs 2° | Male | 0.0268 | 0.0711 | 0.378 | 0.9244 |
| 2° vs 1° | Female | -0.0749 | 0.0473 | -1.583 | 0.2532 |
| 4° vs 1° | Female | -0.0582 | 0.0518 | -1.125 | 0.4983 |

| | | | | | |
|---|---|---|---|---|---|
| 4° vs 2° | Female | 0.0166 | 0.0602 | 0.276 | 0.9588 |

| Epidermal thickness at larval attachment sites | | | | | |
|---|---|---|---|---|---|
| Linear mixed effects regression model: Epidermal thickness ~ infestation number + (1 \| biopsy sample ID) | | | | | |
| Comparisson | Host sex | β | Standard error | T-value | P-value |
| 1° vs uninfested | Male | 12.8 | 10.5 | 1.221 | 0.4624 |
| 4° vs uninfested | Male | 45.3 | 13.3 | 3.405 | 0.0122 |
| 4° vs 1° | Male | 32.5 | 11.6 | 2.801 | 0.0373 |
| 1° vs uninfested | Female | 19.6 | 8.9 | 2.202 | 0.1493 |
| 4° vs uninfested | Female | 70.8 | 10.1 | 7.015 | 0.001 |
| 4° vs 1° | Female | 51.2 | 10.6 | 4.813 | 0.0071 |

| qRT-PCR analysis of leukocytes at tick attachment sites | | | | | |
|---|---|---|---|---|---|
| Linear regression model: Leukocyte marker/β-actin ($Log_{10}$ fold change) ~ infestation number | | | | | |
| Mast cells | | | | | |
| Comparisson | Host sex | β | Standard error | T-value | P-value |
| 1° vs uninfested | Male | 1.007 | 0.233 | 4.33 | 0.0016 |
| 4° vs uninfested | Male | 1.512 | 0.233 | 6.5 | <.0001 |
| 4° vs 1° | Male | 0.505 | 0.233 | 2.17 | 0.1092 |
| 1° vs uninfested | Female | 0.823 | 0.56 | 1.47 | 0.3388 |
| 4° vs uninfested | Female | 1.149 | 0.597 | 1.926 | 0.1738 |
| 4° vs 1° | Female | 0.326 | 0.62 | 0.526 | 0.8601 |
| Eosinophils | | | | | |
| Comparisson | Host sex | β | Standard error | T-value | P-value |
| 1° vs uninfested | Male | 2.7 | 0.301 | 8.966 | <.0001 |
| 4° vs uninfested | Male | 3.668 | 0.301 | 12.18 | <.0001 |
| 4° vs 1° | Male | 0.968 | 0.301 | 3.215 | 0.015 |
| 1° vs uninfested | Female | 1.09 | 0.65 | 1.673 | 0.2548 |
| 4° vs uninfested | Female | 2.45 | 0.693 | 3.53 | 0.0107 |
| 4° vs 1° | Female | 1.36 | 0.721 | 1.886 | 0.185 |
| Neutrophils | | | | | |
| Comparisson | Host sex | β | Standard error | T-value | P-value |
| 1° vs uninfested | Male | 3.06 | 0.3 | 10.2 | <.0001 |
| 4° vs uninfested | Male | 4.19 | 0.3 | 13.954 | <.0001 |
| 4° vs 1° | Male | 1.13 | 0.3 | 3.754 | 0.0051 |
| 1° vs uninfested | Female | 1.391 | 0.694 | 2.005 | 0.1534 |
| 4° vs uninfested | Female | 2.74 | 0.74 | 3.702 | 0.0079 |
| 4° vs 1° | Female | 1.35 | 0.769 | 1.752 | 0.2268 |
| Basophils | | | | | |
| Comparisson | Host sex | β | Standard error | T-value | P-value |
| 1° vs uninfested | Male | 2.36 | 0.272 | 8.676 | <.0001 |
| 4° vs uninfested | Male | 3.45 | 0.272 | 12.683 | <.0001 |
| 4° vs 1° | Male | 1.09 | 0.272 | 4.007 | 0.0031 |
| 1° vs uninfested | Female | 1.4 | 0.594 | 2.358 | 0.0856 |
| 4° vs uninfested | Female | 2.83 | 0.634 | 4.473 | 0.002 |
| 4° vs 1° | Female | 1.43 | 0.658 | 2.176 | 0.1161 |
| T Lymphocytes | | | | | |

| Comparisson | Host sex | β | Standard error | T-value | P-value |
|---|---|---|---|---|---|
| 1° vs uninfested | Male | 1.7 | 0.296 | 5.763 | 0.0001 |
| 4° vs uninfested | Male | 2.82 | 0.296 | 9.549 | <.0001 |
| 4° vs 1° | Male | 1.12 | 0.296 | 3.786 | 0.0048 |
| 1° vs uninfested | Female | 1.23 | 0.666 | 1.852 | 0.1949 |
| 4° vs uninfested | Female | 2.68 | 0.71 | 3.773 | 0.0069 |
| 4° vs 1° | Female | 1.44 | 0.738 | 1.958 | 0.1652 |
| Macrophages | | | | | |
| Comparisson | Host sex | β | Standard error | T-value | P-value |
| 1° vs uninfested | Male | 0.602 | 0.224 | 2.69 | 0.0419 |
| 4° vs uninfested | Male | 0.744 | 0.224 | 3.325 | 0.0121 |
| 4° vs 1° | Male | 0.142 | 0.224 | 0.635 | 0.8033 |
| 1° vs uninfested | Female | 1.85 | 0.537 | 3.447 | 0.0124 |
| 4° vs uninfested | Female | 2.355 | 0.572 | 4.116 | 0.0038 |
| 4° vs 1° | Female | 0.505 | 0.595 | 0.849 | 0.6808 |

| Nucleated cell counts within larval midguts | | | | | |
|---|---|---|---|---|---|
| Negative binomial generalized linear mixed effect model: Nucleated cell count ~ host sex + infestation number + (1 \| larval cohort + infestation date + tick ID) | | | | | |
| Comparisson | Host sex | β | Standard error | T-value | P-value |
| 2° vs 1° | Male | 1.58 | 0.27 | 5.848 | <.0001 |
| 3° vs 1° | Male | 2.61 | 0.263 | 9.923 | <.0001 |
| 4° vs 1° | Male | 2.73 | 0.262 | 10.396 | <.0001 |
| 3° vs 2° | Male | 1.03 | 0.216 | 4.751 | <.0001 |
| 4° vs 2° | Male | 1.15 | 0.216 | 5.319 | <.0001 |
| 4° vs 3° | Male | 0.12 | 0.206 | 0.583 | 0.9372 |
| 2° vs 1° | Female | 0.573 | 0.241 | 2.375 | 0.082 |
| 3° vs 1° | Female | 1.35 | 0.232 | 5.823 | <.0001 |
| 4° vs 1° | Female | 1.597 | 0.234 | 6.816 | <.0001 |
| 3° vs 2° | Female | 0.777 | 0.221 | 3.519 | 0.0024 |
| 4° vs 2° | Female | 1.024 | 0.221 | 4.642 | <.0001 |
| 4° vs 3° | Female | 0.246 | 0.212 | 1.163 | 0.6501 |

| *A.phagocytophilum* burdens in replete larvae | | | | | |
|---|---|---|---|---|---|
| Comparisson | Host sex | β | Standard error | T-value | P-value |
| 5° vs 1° | Male | 0.558 | 0.5615 | 0.9937 | 0.3438 |
| 5° vs 1° | Female | 2.663 | 0.913 | 2.917 | 0.0072 |

| *B. burgdorferi* burdens in replete larvae | | | | | |
|---|---|---|---|---|---|
| Comparisson | Host sex | β | Standard error | T-value | P-value |
| 5° vs 1° | Male | 45.93 | 19.81 | 2.318 | 0.0279 |
| 5° vs 1° | Female | 2.075 | 1.344 | 1.544 | 0.1402 |
