## Supplementary material for "Acquired tick resistance in *Peromyscus leucopus* alters *Ixodes scapularis* infection": Table 3

| Infestation | Sample number | Mouse ID | Mouse sex | Location on body | Histologic lesion | Severity | Epidermal thickness at bite site (μm) | Epidermal thickness away from bite site (μm) |
|---|---|---|---|---|---|---|---|---|
| Not infested | A1 | A | F | Concave pinna | None | N/A | N/A | 18.4 |
| Not infested | A2 | A | F | Convex pinna | None | N/A | N/A | 17.6 |
| Not infested | A3 | A | F | Convex pinna base | None | N/A | N/A | 15.2 |
| Not infested | A4 | A | F | Top of head | None | N/A | N/A | 17.6 |
| Not infested | B1 | B | M | Concave pinna | None | N/A | N/A | 14.4 |
| Not infested | B2 | B | M | Convex pinna | None | N/A | N/A | 13.6 |
| Not infested | B3 | B | M | Convex pinna base | None | N/A | N/A | 9.6 |
| Not infested | B4 | B | M | Top of head | None | N/A | N/A | 12.0 |
| 1° | C1 | C | F | Convex pinna base | Lymphohistiocytic and eosinophilic dermatitis | Mild | 51.2 | 13.6 |
| 1° | C2 | C | F | Convex pinna base | Lymphohistiocytic and eosinophilic dermatitis | Moderate | 38.4 | 19.2 |
| 1° | C3 | C | F | Top of head | Lymphohistiocytic and eosinophilic dermatitis | Mild | 20.8 | 10.4 |
| 1° | D1 | D | M | Convex pinna | Lymphohistiocytic dermatitis | Mild | 31.2 | 16.0 |
| 1° | D2 | D | M | Concave pinna | Lymphohistiocytic and eosinophilic dermatitis | Mild | 42.4 | 15.2 |
| 1° | D3 | D | M | Concave pinna, adjacent to D2 | Lymphohistiocytic dermatitis | Mild | 21.6 | 14.4 |
| 1° | D4 | D | M | Convex pinna base | Lymphohistiocytic and eosinophilic dermatitis | Moderate | 18.4 | 8.8 |
| 1° | E1 | E | M | Convex pinna base | Lymphohistiocytic and eosinophilic dermatitis | Mild | 14.4 | 13.6 |
| 1° | E2 | E | M | Convex pinna | Lymphohistiocytic and eosinophilic dermatitis | Moderate | 20.8 | 12.0 |
| 1° | E3 | E | M | Concave pinna | Lymphocytic dermatitis | Mild | 23.2 | 15.2 |
| 1° | E4 | E | M | Convex pinna base | Lymphohistiocytic and eosinophilic dermatitis | Mild | 30.5 | 8.0 |
| 1° | E5 | E | M | Top of head | Lymphohistiocytic and eosinophilic dermatitis | Moderate | 24.0 | 11.2 |
| 4° | F1 | F | F | Convex pinna base | Histiocytic and eosinophilic dermatitis, cellulitis, and myositis with epidermal ulceration | Severe | 96.8 | 13.6 |
| 4° | F2 | F | F | Convex pinna base, adjacent to F1 | Histiocytic and eosinophilic dermatitis, cellulitis, and myositis with epidermal ulceration | Severe | 79.2 | 14.4 |
| 4° | G1 | G | M | Convex pinna | Lymphohistiocytic and eosinophilic dermatitis and cellulitis with epidermal ulceration | Severe | 35.2 | 16.8 |
| 4° | G2 | G | M | Top of head | Lymphohistiocytic and eosinophilic dermatitis and cellulitis | Severe | 105.0 | 20.0 |
| 4° | G3 | G | M | Convex pinna | Lymphohistiocytic and eosinophilic dermatitis and cellulitis | Severe | 32.8 | 9.6 |
