## Supplementary material for "Acquired tick resistance in *Peromyscus leucopus* alters *Ixodes scapularis* infection": Table 4

**Table 4.** Oligonucleotide primers used in this study

| Name | Target gene | Primer sequence |
|---|---|---|
| *M. musculus* β-actin | XM_030254057.1 | F 5’-ACGCAGAGGGAAATCGTGCGTGAC-3’<br>R 5’-ACGCGGGAGGAAGAGGATGCGGCAGTG-3’ |
| *P. leucopus* eosinophil major basic protein (*embp*) | XM_028894239.2 | F 5’-GGAGATATGAAGCAGCCCCT-3’<br>R 5’-GGCCTCCAGATGAAAAGCAG-3' |
| *P. leucopus* neutrophil myeloperoxidase (*mpo*) | XM_028878233.2 | F 5’-ATTGCATTCCCTTCTTCCGC-3’<br>R 5’-GGTCCTCGCTGCCATATACT-3’ |
| *P. leucopus* mast cell protease 4 (*mcpt4*) | XM_028892202.2 | F 5’-GTGATAACGGCTGCACACTG-3’<br>R 5’-GTGTGGGCTCTTTCTTGCTC-3’ |
| *P. leucopus* basophil granzyme-like protein 2 (*mcpt8*) | XM_028892261.2 | F 5’-GGTACAGAGTCCAAACCCCA-3’<br>R 5’-TCTCTCACCAGGAAACCACC-3’ |
| *P. leucopus* macrophage allograft inflammatory factor 1 (*iba1*) | XM_037200022.1 | F 5’-CCATCTCCCCACCTAAGACC-3’<br>R 5’-GCTTTTCCTCCCTGCAAGTC-3’ |
| *P. leucopus* T lymphocyte CD3 ε chain (*cd3*) | XM_037207548.1 | F 5’-AGGCCCAGATTTCCTCAGAC-3’<br>R 5’-CGTCACTTGGCAATGTTCGA-3’ |
| *I. scapularis* actin | XM_029977298.1 | F 5’-GCCGGGACCTTACAGACTATC-3’<br>R 5’-CACGGACAATTTCACGCTCG-3’ |
| *B. burgdorferi flaB* | MN954474.1 | F 5’-TTGCTGATCAAGCTCAATATAACCA-3’<br>R 5’-TTGAGACCCTGAAAGTGATGC-3’ |
| *A. phagocytophilum 16S* | NC_007797 | F 5’-CCCTAAGGCCTTCCTCACTC-3’<br>R 5’-CAGCCACACTGGAACTGAGA-3’ |
